# Cultivation of Exoelectrogenic Bacteria in Conductive DNA Nanocomposite Hydrogels Yields a Programmable Biohybrid Materials System

**DOI:** 10.1101/864967

**Authors:** Yong Hu, David Rehnlund, Edina Klein, Johannes Gescher, Christof M. Niemeyer

**Affiliations:** Karlsruhe Institute of Technology (KIT), Institute for Biological Interfaces (IBG 1), Hermann-von-Helmholtz-Platz 1, 76344, Eggenstein-Leopoldshafen, Germany; Karlsruhe Institute of Technology (KIT), Institute for Applied Biosciences (IAB), Fritz-Haber-Weg 2, 76131, Karlsruhe, Germany

**Keywords:** Carbon nanotubes, DNA, silica nanoparticles, nanocomposites, rolling circle amplification, *Shewanella*, extracellular electron transfer

## Abstract

The use of living microorganisms integrated within electrochemical devices is an expanding field of research, with applications in microbial fuel cells, microbial biosensors or bioreactors. We describe the use of porous nanocomposite materials prepared by DNA polymerization of carbon nanotubes (CNT) and silica nanoparticles (SiNP) for the construction of a programmable biohybrid system containing the exoelectrogenic bacterium *Shewanella oneidensis*. We initially demonstrate the electrical conductivity of the CNT-containing DNA composite by employment of chronopotentiometry, electrochemical impedance spectroscopy, and cyclic voltammetry. Cultivation of *Shewanella oneidensis* in these materials shows that the exoelectrogenic bacteria populate the matrix of the composite, while non-exoelectrogenic *Escherichia coli* remain on its surface. Moreover, the ability to use extracellular electron transfer pathways is positively correlated with number of cells within the conductive synthetic biofilm matrix. The *Shewanella* containing composite remains stable for several days. Programmability of this biohybrid material system is demonstrated by on-demand release and degradation induced by a short-term enzymatic stimulus. The perspectives of this approach for technical applications are being discussed.

## 1. Introduction

Electrically conductive composite biomaterials that combine the redox switching and electrical properties of inherently conductive components with the efficient small molecule transport, high hydration values and biocompatibility of cross-linked hydrogels are considered promising materials for applications in eukaryotic cell systems and higher organisms, such as drug release devices or implantable electronics.^1^ In addition to the importance of conductive materials for biomedical applications, the electrically conductive connection of technical and living systems also plays an important role in biotechnology.^2^ For example, the integration of living microorganisms into bioelectrochemical devices is driving novel technical applications ranging from microbial biosensors^3-4^ and fuel cells^5-6^ to novel bioreactors.^7^ A central point of these biotechnological developments are so-called exoelectrogenic bacteria, which are able to link their metabolism with redox processes on electrodes. Electron transfer can occur by direct physical contact between electrodes and outer membrane cytochromes and/or conductive cellular appendages of bacteria, or by rapid indirect electron transport via endogenous mediators produced by some bacteria.^8-11^

Considerable efforts have been made to improve the electron transfer rate at the bacteria-electrode interface by coating the electrodes with micro- and nanostructured conductive materials.^12-16^ For example, a conductive carbon nanotube (CNT) hydrogel prepared by electrodeposition of CNT and chitosan onto carbon paper electrodes was used for enrichment of electrochemically active bacteria from anaerobic sludge.^15^ Although improved conductivity has been observed in these studies, the materials described are generally limited to serving as a 2D interface for the binding of microorganisms. However, in order to comprehensively improve the interface between electronics and biology, hierarchically structured porous 3D scaffolds are required, which have both an open structure for efficient diffusion of nutrients and immigration of large numbers of cells, as well as a large effective contact area with which the microscale electrode elements can be connected efficiently with cellular components on the nanometer length scale. The intimate hybridization of such materials with living microorganisms could open the door to novel biohybrid material systems that efficiently exploit optimized cell-material interactions for biotechnological processes.

Based on recent work on the application of porous nanocomposite materials accessible by DNA polymerization of CNT and silica nanoparticles (SiNP) as bioinductive coatings for eukaryotic cells,^17^ we report here on the use of these materials for the construction of a programmable biohybrid system. To this end, we initially demonstrate electrical conductivity of the CNT-containing composite by employment of chronopotentiometry (CP), electrochemical impedance spectroscopy (EIS), and cyclic voltammetry (CV). Studies on the cultivation of *Shewanella oneidensis* wild type and mutants in key genes for extracellular electron transfer reveal that the ability to conduct electron transfer to the cell surface is key for a successful growth within the material. This is also corroborated by the very limited growth observed for the non-exoelectrogenic organism *Escherichia coli*. The *Shewanella* containing composite remains stable for several days and shows electrochemical activity. Programmability of this biohybrid material system is demonstrated by on-demand release and degradation induced by a short-term enzymatic stimulus. The perspectives of this approach for technical applications are being discussed.

## 2. Materials and methods

### Synthesis of SiNP/CNT-DNA nanocomposite materials

Synthesis of SiNP, chemical conjugation of SiNP and CNT with DNA oligonucleotide primers, ligation of rolling circle amplification (RCA) template (T) on SiNP-primer (CNT-P) and CNT-primer (CNT-P) and RCA polymerization of the nanoparticles were performed as previously described.^17^ In brief, linear ssDNA (T, 10 µM, 30 µL) and 10X T4 DNA ligation buffer (500 mM Tris-HCl, 100 mM MgCl_2_, 10 mM ATP, 100 mM dithiothreitol (DTT), 7.5 µL) were added to 60 μL SiNP-P suspension (10 mg/mL) or CNT-P-T (800 µg/mL), followed by addition of T4 DNA ligase (400,000 U/mL, 2.5 µL, New England Biolabs) to ligate the nicked ends of the T oligonucleotide at 25 °C for 3 h. To synthesize SiNP/CNT-DNA hydrogels, the so-produced SiNP-P-T (50 µL) or CNT-P-T (50 µL) were then polymerized via RCA by mixing with dNTPs (10 mM, 10 µL), 10X BSA (10 mg/mL, 5 µL), 10X phi29 DNA polymerase buffer (500 mM Tris-HCl, 100 mM MgCl_2_, 100 mM (NH_4_)_2_SO_4_, 40 mM DTT, pH 7.5, 5 µL) and phi29 DNA polymerase (10,000 U/mL, 5 µL, New England Biolabs) at 30 °C for 48 h. Additional details on chemicals and oligonucleotide sequences (Table S1) are specified in the Supplementary Information.

### Materials characterization and bacteria cultivation

All experimental details concerning electrochemical measurements, scanning electron microscopy (SEM) analysis, fluorescence imaging analysis and bacteria cultivation methods are provided in the Supplementary Information.

## 3. Results

As a potential candidate for an improved interface between electronics and microbiology, we chose a hierarchically structured porous 3D nanocomposite hydrogel comprised of SiNP, CNT and DNA strands. As previously reported^17^, this material is produced from DNA-coated SiNP and CNT particles that are interwoven with each other by enzymatic polymerization using the Phi29 DNA polymerase-catalyzed rolling circle amplification (RCA, see Fig 1a). For this purpose, SiNP (approx. 80 nm in diameter) and CNT (1 μm in length and 0.83 nm in diameter) coated with RCA primers were allowed to hybridize with a linear ssDNA oligomer (T) that was subjected to ring closure by enzymatic ligation. The resulting particles then served as the template for RCA. Three different materials were produced this way, namely binary composites that contained only SiNP or CNT (in the following denoted as *S* and *C*, respectively) or the ternary composite containing both SiNP and CNT, denoted as *SC*.

**Figure 1.**
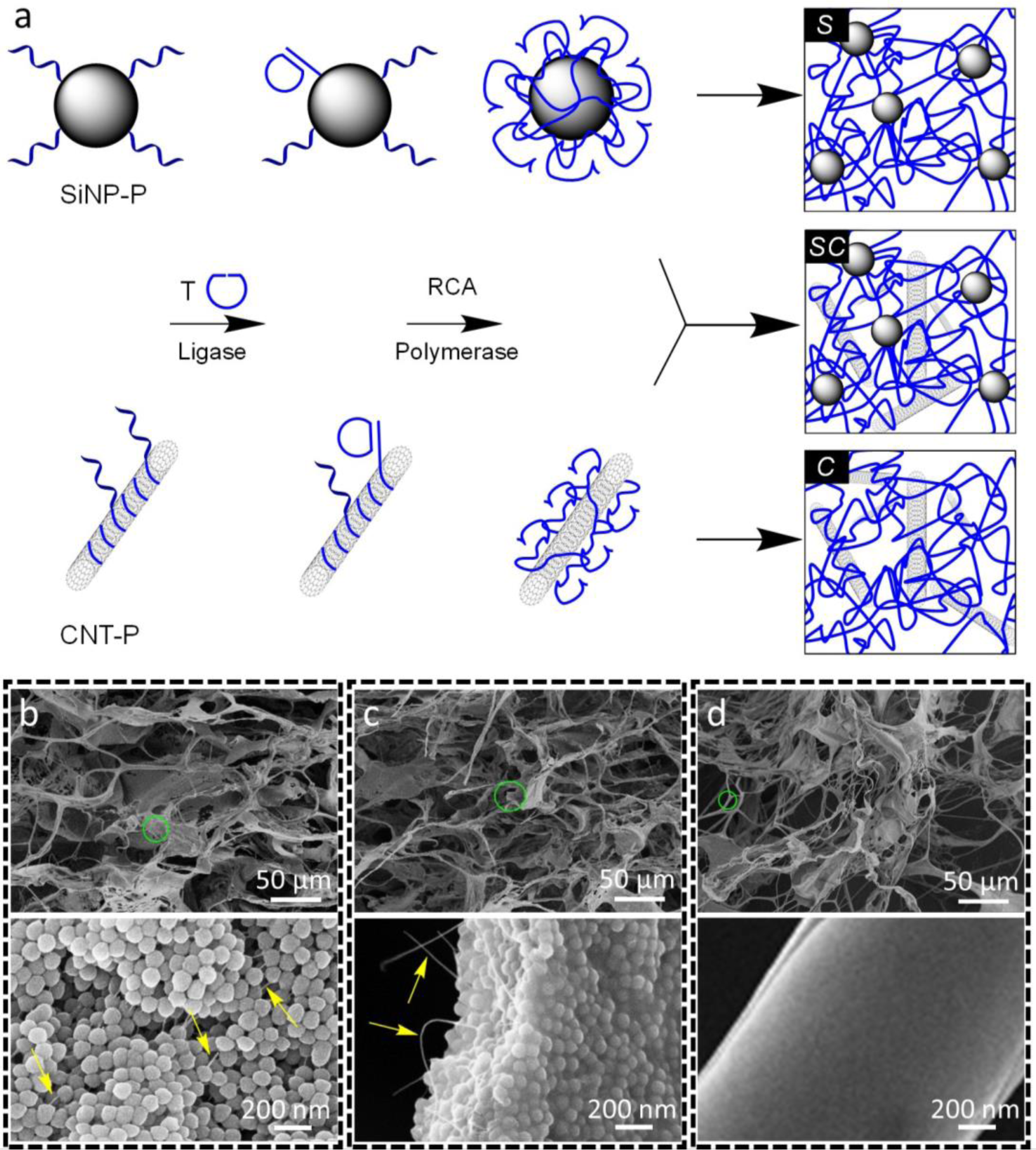
Synthesis of binary and ternary SiNP/CNT-DNA nanocomposite materials. (**a**) Schematic illustration of RCA-based synthesis. DNA oligonucleotide primer modified SiNP-P and/or CNT-P are used for enzymatic cyclisation of RCA template T and subsequent polymerization to yield the corresponding binary (*S* or *C*) or ternary (*SC*) DNA nanocomposites. (**b**-**d**) Representative SEM images of *S, C*, and *SC* materials at different magnifications. Magnifications of the green circled areas in **b** - **d** are shown in the images below. The arrows point at DNA polymers in (**b)** and CNT in (**c)**, respectively. Note that individual CNT were not resolvable in the *C* materials (**d)**.

The structure and composition of the hydrogels was characterized by scanning electron microscopy (SEM). The *S* materials have an amorphous morphology with a distinctive hierarchical ultrastructure, which clearly reveals the DNA-coated nanoparticles that are connected by DNA filaments (Fig. 1b). The ternary *SC* and the binary *C* composite materials showed a similar mesoscopic morphology (Fig. 1c, d). Individual CNT could be observed in the *SC* composite materials (Fig. 1c) but were not resolvable in the *C* materials (Fig. 1d). Hence, the presence of CNT in the *C* materials had been confirmed by Raman microscopy.^17^

The electronic and electrochemical properties of SiNP/DNA-CNT nanocomposite materials in both hydrated and dehydrated states were studied in detail by using chronopotentiometry (CP), electrochemical impedance spectroscopy (EIS), and cyclic voltammetry (CV), respectively. CP and EIS analyses of the nanocomposite materials provided insights into the conductivity of the materials under constant and frequency dynamic conditions, respectively (Fig. 2). Conductivity measurements were performed on screen printed electrodes with interdigitated arrays (IDA) of gold lines with varying distances (i.e., 10, 100 and 200 µm) between the working and counter electrode lines (Fig. 2, see also Tables S2 and S3). In hydrated state, CP analysis revealed that the *SC* materials displayed higher conductivity (11-21 µS/m) than the *S* materials (2-8 µS/m). It was also found that the SiNP did not lower the conductivity of *SC* materials when compared with *C* materials (3-20 µS/m) (Fig. 2b). The EIS data were fitted to an equivalent circuit (Fig. 2a) and the resulting Bode plots showed a constant impedance in the 1 MHz to 1 kHz range during which the phase dropped towards 0° (Fig. 2c). At frequencies below 1 kHz, a steady increase in impedance was observed, which was coupled to the increase tendency in the wave phase to 60-80°. This phase fluctuating impedance behavior indicates a transition from resistive to capacitive dominated circuit, since 0° and -90° correspond to pure resistance and capacitance behavior, respectively.^18^

**Figure 2.**
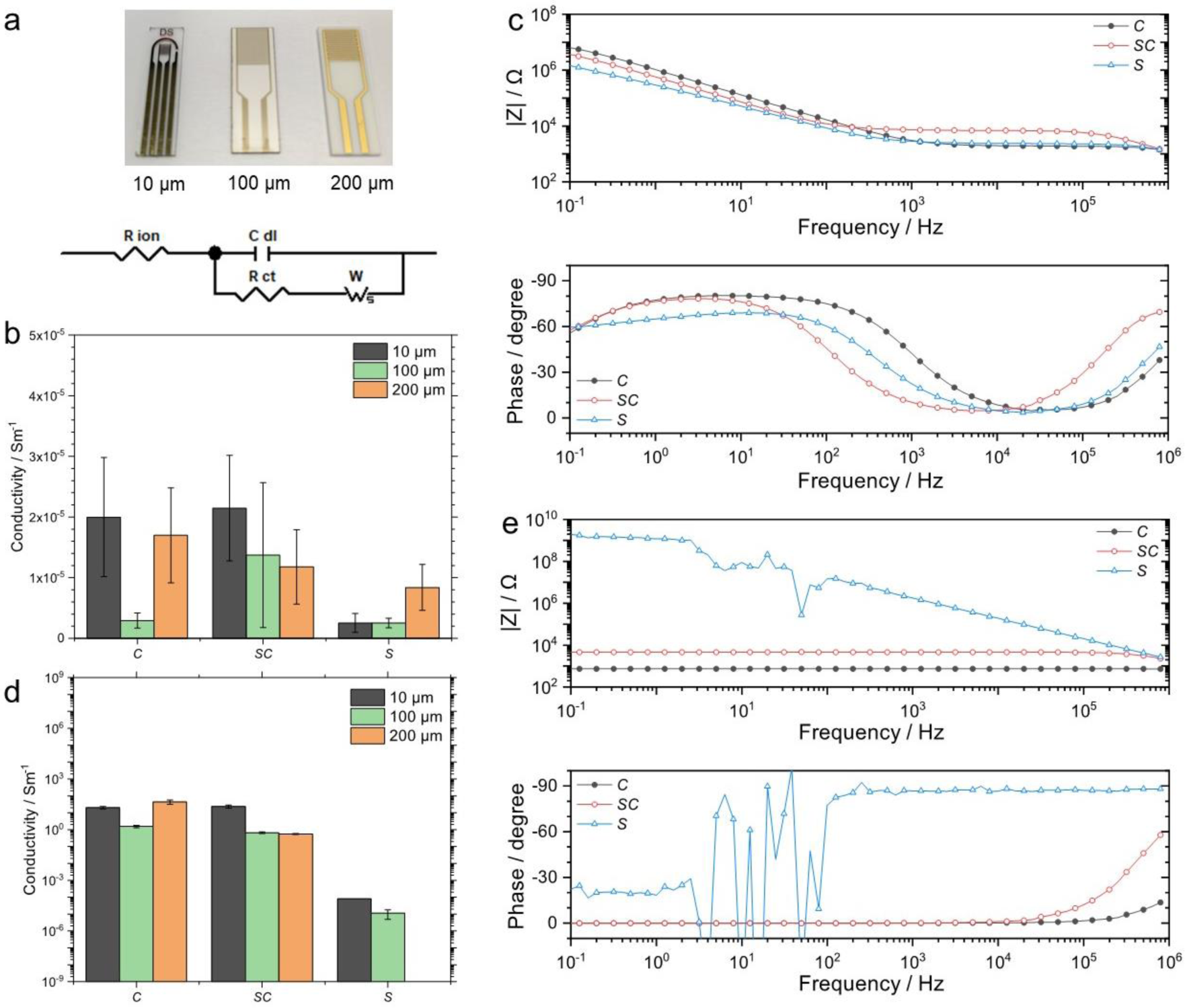
Characterization of electrical properties of SiNP/CNT-DNA nanocomposite materials. (**a**) Photo of IDA electrodes used to investigate the electrical conductivity and equivalent circuit model used to fit the EIS data. Electrical properties of nanocomposite materials investigated under (**b, d**) direct current and (**c, e**) alternating current. Electrical conductivity investigation of nanocomposite materials (*S, SC* and *C*) in (**b, c**) hydrated and (**d, e**) dehydrated states. (**c, e**) Bode plots illustrate the impedance modulus and phase dependence on the frequency in the 0.1 Hz to 1 MHz range using a 100 µm IDA electrode.

In dehydrated state, CP analysis revealed that both the *C* materials (1.6-43 S/m) and *SC* materials (0.6-23 S/m) have a similar conductivity, which is several orders of magnitude higher than that of the *S* materials (11-79 µS/m) (Fig. 2d). EIS analysis of the dehydrated nanocomposite materials revealed that the dehydrated *C* and *SC* materials exhibit a constant impedance throughout the entire frequency spectrum, whereas the *S* materials showed a continuous increase in impedance towards lower frequencies (Fig. 2e). Phase analysis indicated that, apart from an initial charging of the double layer at high frequencies, the CNT containing materials exhibit pure resistive behavior in contrast to the *S* materials which showed close to pure capacitive behavior in the majority of the frequency range (Fig. 2e). These results indicate that the CNT network offers electronic conduction for *C* and *SC* materials while the *S* materials behave like an insulator in their dehydrated state.

Nyquist plots show a similar behavior for all nanocomposite materials in the hydrated state with an initial well-defined semicircle followed by a linear increase in impedance towards lower frequencies (Fig. S1, Supplementary Information). This impedance behavior is indicative of an initial charging of the double layer followed by a Warburg impedance for the ionic movement in the hydrogel matrix. With no clear distinction between the *S* materials and *C*/*SC* materials, it was reasonable to assume that the Warburg impedance is controlled by ionic movement along the DNA backbone. However, after dehydration, it seemed the material had collapsed into an initial double layer charging followed by pure electronic conduction with a constant resistance of 0.68 kΩ and 2.2 kΩ for the *C* and *SC* materials, respectively.

Altogether, the electrical conductivity properties of the *C* and *SC* materials are comparable to other conductive polymer-based hydrogels.^1, 19-21^ The DNA backbone in the *C* and *SC* materials appears to inherit the ionic conductive property observed in conventional conductive polymers. While it is known that the CNT additives offer an excellent local electrical conductivity,^22-23^ full activation of the CNT electrical network cannot be seen in the hydrated *C* and *SC* materials over 10-200 µm distances, presumably because the length of CNT (approx. 1 µm) is shorter than the gap distance and the CNT particles are not sufficiently aligned and connected. However, the complete supported conduction of CNT is observed in the dehydrated state, thus suggesting that the collapse of the material structure brings the CNT in sufficiently close contact.

While the above results suggested that the nanocomposite materials could be useful for bioelectrochemical systems comprising living bacteria, it was also necessary to investigate whether the materials exhibit the electrochemical stability in the desired potential range. Therefore, CV measurements were performed to investigate the electrochemical activity of the nanocomposite materials, in the absence of a redox mediator (Fig. S2). We observed only one clear faradaic peak for all three nanocomposite materials at potentials below -0.8 V (vs. Ag/AgCl) corresponding to hydrogen gas evolution. Interestingly, the binary *S* and *C* materials were observed to influence the magnitude of the hydrogen reaction, whereas, surprisingly, the ternary *SC* materials showed the slowest kinetics. Importantly, no faradaic reactions were observed in the potential range -0.4 to 0 V, which is the most relevant range for the majority of bioelectrochemical systems. Hence, these results confirmed that the DNA backbone is electrochemically stable and the composites are suitable for use in bioelectrochemical applications.

We then investigated the utility of SiNP/CNT-DNA nanocomposite materials as substrata for cultivation of bacteria. To this end, both *E. coli* and *S. oneidensis* were allowed to grow for 24 h in LB medium inside microplate wells coated with a 45-50 μm layer of the DNA nanocomposite materials. Analysis by confocal laser scanning microscopy (CLSM, Fig. 3, see also Figs. S3, S4) revealed that *E. coli* formed a biofilm on top of the *S* material (Fig. 3a), whereas the bacteria did not well proliferate in wells coated with *SC* materials (Fig. 3b). We attribute the reduced growth of *E. coli* to the previously observed toxicity of CNT to these bacteria.^24-26^

**Figure 3.**
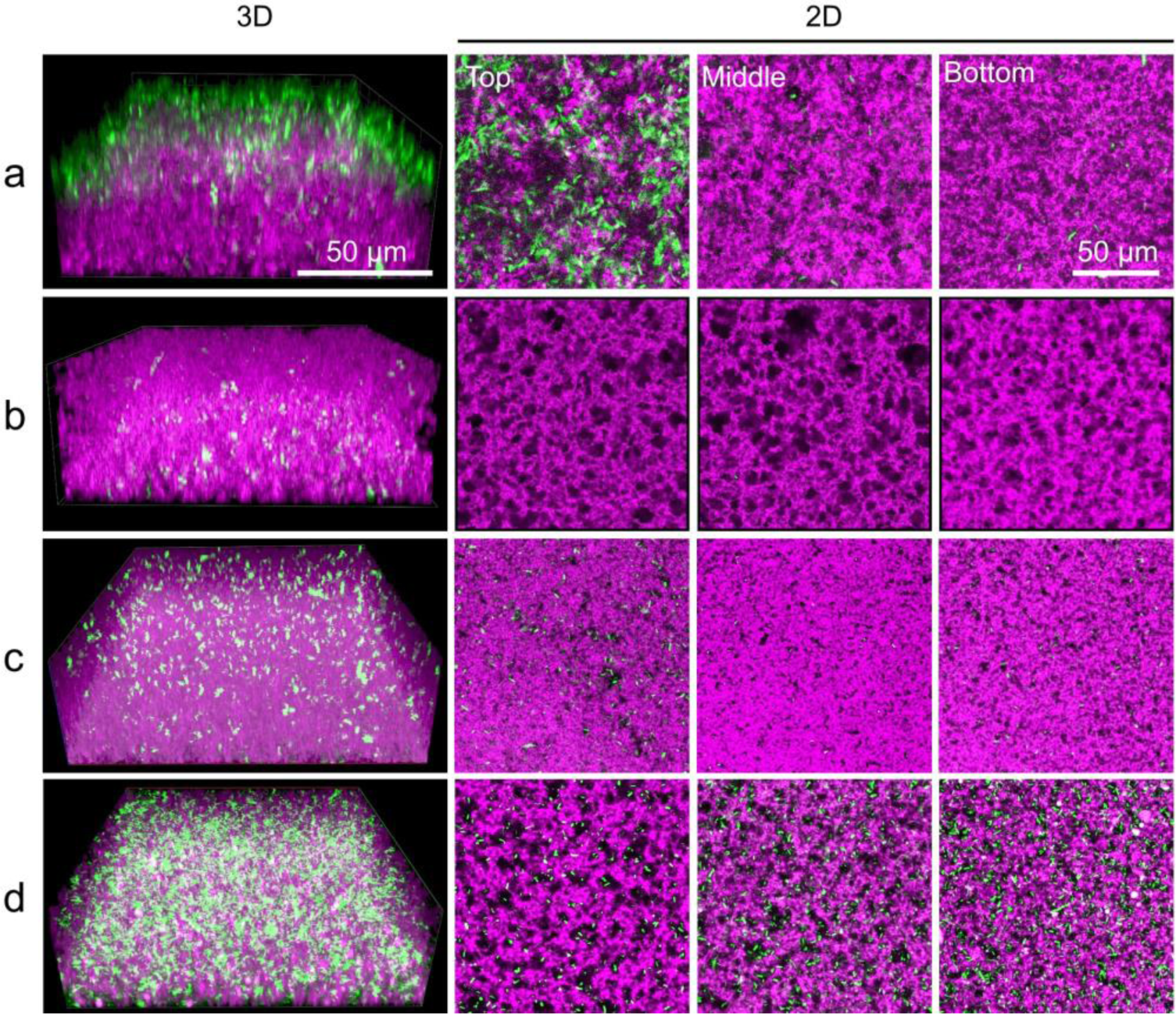
Cultivation of bacteria in SiNP/CNT-DNA nanocomposite materials. Representative confocal fluorescence images of (**a, b**) *E. coli* and (**c, d**) *S. oneidensis* grown in *S* (**a, c**) and *SC* (**b, d**) composite materials in LB medium. Images were acquired after 24 h of growth.

In contrast, *S. oneidensis* showed a completely different behavior. These exoelectrogenic bacteria proliferated only weakly on top of the *S* material (Fig. 3c) while a strong growth occurred within the *SC* material (Fig. 3d, see also Fig. S4). The observation that the presence of CNT in the *SC* material promotes the proliferation of *S. oneidensis* can be explained by the fact that CNT can transfer and dissipate the metabolic electrons from the cells throughout the nanocomposite, thereby overcompensating any negative toxic side effects to the cells. This hypothesis is in agreement with the earlier observation that CNT can act as electrode modifier to promote the direct electron transfer from *S. oneidensis*.^13^ The distinctive differences of the growth of *S. oneidensis* on either *S* or *SC* materials was also confirmed by SEM analysis, where the formation of densely grown bacteria populations on *SC* material was clearly evident (Fig. S5).

To further elucidate the hypothesis that the CNT play a role in electron exchange processes between the bacteria and the materials, quantitative analyses of cell growth were conducted with *S. oneidensis* wild type and two *S. oneidensis* mutants. The mutants are deficient either in all outer membrane cytochrome encoding genes (*Δomc*)^27^ or only the gene for the inner membrane menaquinol-oxidase CymA (*ΔcymA*).^28^ Since the minimal medium lacks potential electron acceptors and electron shuttles, bacterial metabolism and growth depend on whether the metabolic electrons of the cells can be efficiently dissipated into the material matrix and finally to oxygen as the only available terminal electron acceptor of the system. It was expected that the diffusion of oxygen into the material would be growth limiting. Hence, a transfer of respiratory electrons via an interplay of CNT, outer membrane cytochromes and endogenous electron shuttles would support growth, as these structures would bring respiratory electrons in the vicinity of the material and would trigger a cell free oxygen reduction. Both mutants lack key elements of extracellular electron transfer. CymA is involved in the electron transport to all other anaerobic electron acceptors that *S. oneidensis* can use. In contrary, the *Δ omc* strain has a more narrow phenotype as it is only unable or severely affected to respire with metals or anodes as electron acceptors.^29^ Hence, we hypothesized that both mutant strains should have a clear growth disadvantage compared to the wild type.

To test on this hypothesis, the three bacterial strains were seeded in wells of a microtiterplate bearing a layer of *SC* material, cultivated in minimal medium and analyzed by CLSM. Since the bacterial growth in minimal medium proceeds more slowly than in LB medium, the cells were cultured for a longer time period (48 h) as compared to the culturing in LB medium. Similar as in full LB medium, the *S. oneidensis* wild type completely penetrated the *SC* material and dense cell populations were preferentially found in bottom regions of the materials layer (Figs. 4a, b, see also Figs. S6, S7). As expected, the lack of both the outer membrane cytochromes or CymA, respectively, led to a significant reduction in the propagation capability, with the *Δomc* strain being affected to a lesser extent than the *ΔcymA* strain. The results also suggested that the conductive *SC* materials can compensate better for the negative effects of the absence of the outer membrane cytochromes (*Δomc* strain) than for the absence of CymA (*ΔcymA* strain). Altogether the data fully support our hypothesis that the metabolic electrons from the cells can be transferred and dissipated by the surrounding nanocomposite materials. Given the different growth rates of the *Δomc* and *ΔcymA* strains, it appears that the conductive nanocomposite materials may be in direct physical contact with the cell surface to play a similar role as the terminal reductases, while no interactions occur between the materials and the cytoplasmic membrane.

**Figure 4.**
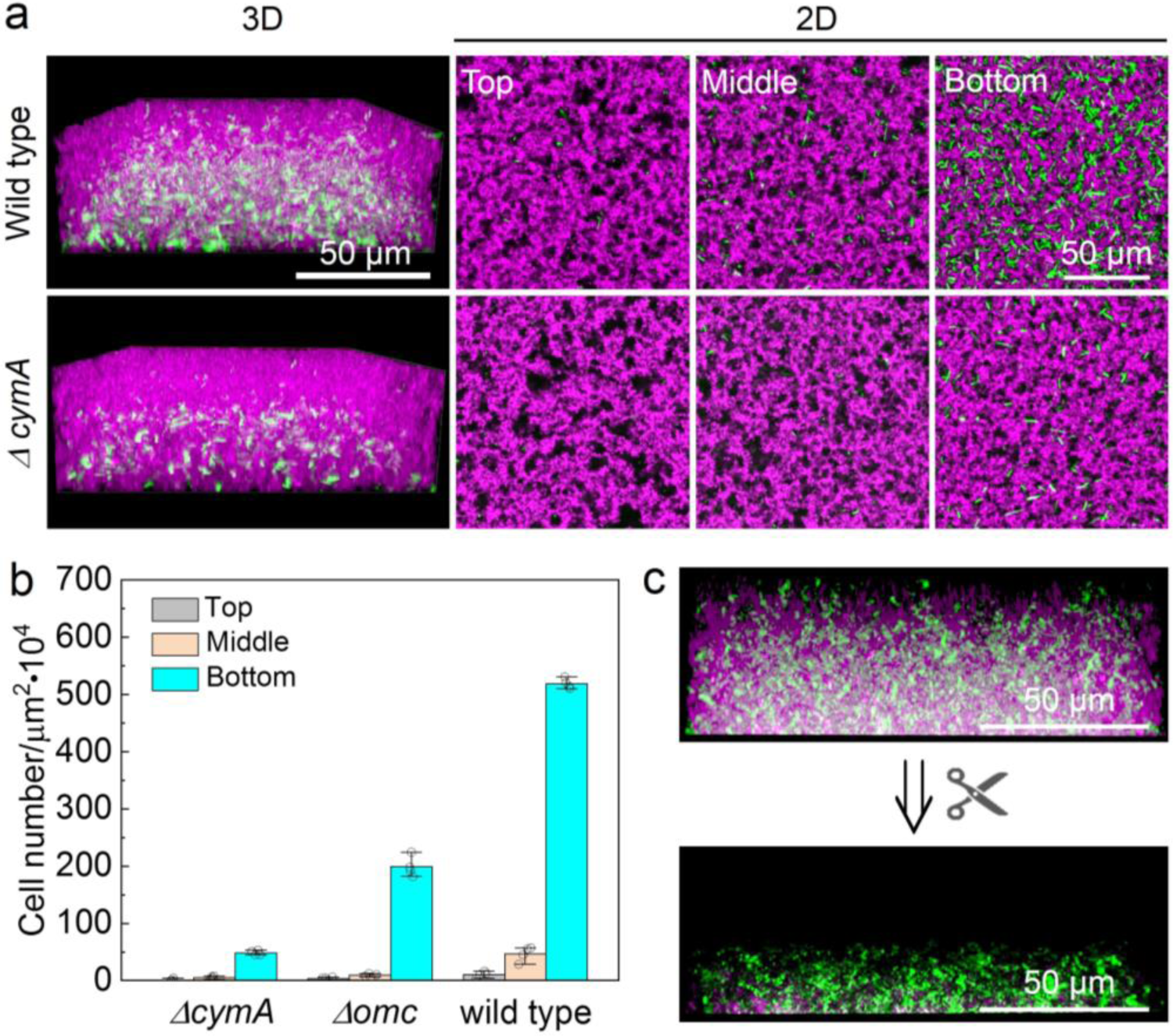
Bacterial propagation in *SC* composite and enzymatic release of the resulting biohybrid material. (**a**) Fluorescence imaging analysis of *S. oneidensis and S. oneidensis Δ cymA* grown in *SC* nanocomposite materials in minimal medium for 48 h. For representative images of all strains, see Figs. S6, S7. (**b**) Statistical analysis of cell densities observed in three different regions of *SC* materials determined after 48 h. (**c**) Fluorescence imaging analysis of *S. oneidensis* grown in *SC* material for 24 h (upper panel) and released by enzymatic cleavage of the DNA backbone with BstEII-HF endonuclease (lower panel).

Finally, we wanted to show that bacteria that have grown into DNA nanocomposites can be handled like a programmable material system. For this purpose, a *SC* material-supported *S. oneidensis* biofilm was grown, which led to an interwoven tissue of bacteria and composite material, as shown in the top panel of Fig. 4c. Since the polymer backbone of the *SC* material was designed to possess enzymatic restriction sites specifically addressable with the endonuclease BstEII-HF (Fig. S8), the addition of this enzyme induced the spontaneous degradation of the material system within 1 h (bottom panel, in Fig. 4c). Importantly, the released bacteria could be regrown under standard conditions, thus indicating that that the collected bacteria were not harmed by grow and degradation of the biohybrid material (Fig. S9). Furthermore, control studies with the non-complementary SexAI endonuclease revealed that the biomaterial system remains stable under these conditions (Fig. S10). These results thus show that the self-organization of *S. oneidensis* in SiNP/CNT-DNA nanocomposite materials described here leads to an integrated biohybrid material system.

## 4. Discussion

In summary, we here describe the use of SiNP/CNT-DNA nanocomposite materials as a platform for cultivation of exoelectrogenic bacteria. In previous studies we could show that the mechanical properties of these materials can be adjusted by variation of the CNT and SiNP composition in order to influence the adhesion, migration and proliferation of eukaryotic cell systems.^17^ The studies shown here go beyond a mechanical influence on cellular processes and show, for the first time, that the nanocomposite materials have an electrical conductivity that can be exploited for the cultivation of exoelectrogenic bacteria. Although the exact molecular processes still have to be elucidated in detail, the results shown here provide a conclusive picture that the electrical conductivity of the materials is decisive for the efficient growth of *S. oneidensis*.

The resulting biohybrid material system shows both the properties of the contained biological (metabolism, growth) and the synthetic material components. With respect to the latter, in addition to the mechanical support function and conductivity, the biochemical addressability, is particularly noteworthy, which was examplified by specific enzymatic degradation. This programmability of the polymer backbone, which can be tailored by targeted sequence design over several size scales from the lower nanometer to the upper micrometer range, offers unique advantages over synthetic polymers and thus far-reaching perspectives for the use of DNA material systems to control living organisms.^2, 30^

With regard to the exoelectrogenic bacteria investigated here, possible applications go beyond the currently intensively researched biosensor and fuel cell systems. For example, the cultivation of biofilms in microfluidic systems is making substantial progress^31^ and this technological platform is also being used for research into supramolecular and dissipative material systems.^32-34^ We therefore believe that future advancements of fluidically controllable DNA materials systems can make an important contribution to the development of novel biotechnological production systems.

## Acknowledgements

This work was supported by the Helmholtz programme “BioInterfaces in Technology and Medicine”. Y.H. is grateful to the China Scholarship Council (CSC) for a Ph.D. fellowship. D.R. would like to acknowledge the financial support from The Swedish Research Council (VR-2017-06320). We also thank Volker Zibat (LEM, KIT) for scanning electron microscopy analyses.

## Conflicts of interest

The authors declare that a patent application for SiNP/CNT-DNA composite materials has been filed. Y.H. and C.M.N. are included on the patent and declare competing interest.

## Appendix A. Supplementary data

Supplementary data to this article can be found online at https://doi.org/xxx.

