## Supplemental files for "Cultivation of Exoelectrogenic Bacteria in Conductive DNA Nanocomposite Hydrogels Yields a Programmable Biohybrid Materials System"

### Synthesis of SiNP/CNT-DNA nanocomposite materials

The SiNP/CNT-DNA nanocomposite materials were prepared as previously described.[1] In brief, linear ssDNA oligonucleotdes (T, for all oligonucleotide sequences, see Table S1) were circularized through hybridization with P1 attached on the surface of primer-modified silica nanoparticles (SiNP-P) using T4 DNA ligase. To this end, linear ssDNA (T, 10 µM, 30 µL) and 10× T4 DNA ligation buffer (500 mM Tris-HCl, 100 mM $MgCl_2$, 10 mM ATP, 100 mM dithiothreitol (DTT), 7.5 µL) were added to 60 µL SiNP-P suspension (10 mg/mL), and the mixture was incubated at 25 °C for 3 h. After addition of 2.5 µL T4 DNA ligase (400,000 U/mL, New England Biolabs), the mixture was further incubated for more than 3 h at 25 °C to ligate the nicked ends of the template, leading to the formation of particle-primer-template (SiNP-P-T) complexes. In a similar way, primer-modified carbon nanotubes (CNT-P) were used for preparation of the CNT-P-T (30 µL of T (10 µM) and 60 µL of CNT-P (800 µg/mL).

The RCA reaction mixture contained dNTPs (10 mM, 10 µL), 10× BSA (10 mg/mL, 5 µL), 10× phi29 DNA polymerase buffer (500 mM Tris-HCl, 100 Mm $MgCl_2$, 100 mM $(NH_4)_2SO_4$, 40 mM DTT, pH 7.5, 5 µL) and phi29 DNA polymerase (10,000 U/mL, 5 µL, New England Biolabs). The polymerisation was initiated *via* the addition of 50 µL of the SiNP-P-T or CNT-P-T particles or else the mixture of SiNP-P-T/CNT-P-T. After incubation at 30 °C for 48 h, the SiNP-DNA nanocomposite hydrogels, CNT-DNA nanocomposite hydrogels or SiNP/CNT-DNA nanocomposite hydrogels were purified by carefully replacing the reaction buffer with DPBS for 5-7 times and the nanocomposite hydrogels were collected and stored at 4 °C before use. The so formed SiNP-DNA nanocomposite hydrogel, denoted as *S* nanocomposite material, contained a final SiNP-P concentration of 4 mg/mL, the so formed CNT-DNA nanocomposite hydrogel (*C* nanocomposite material) contained a final CNT-P concentration of 320 µg/mL, and the SiNP/CNT-DNA nanocomposite hydrogel (*SC* nanocomposite material) contained 4 mg/mL SiNP-P and 320 µg/mL of CNT-P. Unless otherwise stated, Cy5@SiNP were used to enable visualization of nanocomposite materials during fluorescence imaging.

### Electrochemical measurements

The electrical properties of SiNP/CNT-DNA nanocomposite materials were characterized by chronopotentiometry (CP), electrochemical impedance spectroscopy (EIS) and cyclic voltammetry (CV) using a potentiostat/galvanostat with built-in impedance module (Gamry, USA). Screen printed interdigitated arrays (IDA) of gold lines with gap/line thicknesses of 10, 100 and 200 µm (Dropsens, Spain) were used electrodes to study the electrical conductivity of the nanocomposite materials. A three-electrode setup was used for the CV analyses with a gold IDA electrode (PINE Research, USA) serving as working and counter electrode in connection to an external Ag/AgCl reference electrode (BASi, USA). The nanocomposite materials were first washed and resuspended in Milli-Q water for the CV measurements. Starting at the open circuit potential (OCP) the potential was swept with a scan rate of 100 mV/s between -1 V and 0.5 V for three cycles. Comparative voltammograms presented in Fig. S2 show cycle 2 for all samples.

Conductivity measurements were performed in both direct current (i.e., chronopotentiometry) and alternating current (i.e., electrochemical impedance spectroscopy) mode. A two-electrode setup was used with the gold IDA electrodes analyzed in Milli-Q water. EIS was first implemented by applying an alternating current sinusoidal signal of 10 mV amplitude versus the OCP. The value of the impedance was determined at 10 frequencies per decade over the range of 0.1 Hz to 1 MHz.

Chronopotentiometric analyses of the direct current conductivity was investigated by applying increasing constant currents of: 10 nA; 100 nA; 500 nA; 1 µA; 5 µA; 10 µA; 50 µA. Each current was applied for 60 s to establish a steady state. Triplicate measurements of each material were performed for the three different IDA electrodes with varying gap lengths. Each hydrogel sample was analyzed in its pristine 'hydrated' form and in its 'dehydrated' form after 24 h drying in room temperature. The conductivity was calculated using Equation 1 and Table S2. The hydrogel thickness was measured by scanning electron microscopy (SEM) in the dried state and by a digital caliper in the wet state. The length was determined by the precise gap distance between each gold line on the IDA electrode. AC resistance values were extracted by fitting the impedance data to an equivalent circuit (Fig. 2a) using the Zview software. The AC resistance is composed of both the charge transfer impedance and the Warburg impedance. Conductivity (S/m) as a function of the resistance (R), area (A) and length (l) of the conductor is presented as Equation 1:

$$\sigma = \frac{RA}{l}$$

### Cultivation of bacteria in the nanocomposite materials

*E. coli* or *S. oneidensis* cells were transformed with a pAra-RFP plasmid to enable the expression of Red Fluorescent Protein (RFP) upon exposure to arabinose containing medium. The arabinose regulatory element pAra was amplified by PCR using the pTF16 vector (Takara Bio Inc.) as a template and inserted into a lab stock pMK vector (GeneArt, Thermo Scientific, containing the RFP variant mRFP1) by Gibson isothermal assembly.[2] The rfp gene was encoded on a pMK-RQ plasmid carrying also the kanamycin resistance gene. The vector was transformed into electrocompetent *E. coli* or *S. oneidensis* cells. Successful cloning was verified by commercial sequencing (LGC genomics).

The applicability of the composite materials for bacterial culture was demonstrated by the use of mRFP1-expressing *E. coli* and *S. oneidensis* wild type. To investigate the impact of extracellular electron transfer processes on the bacterial growth within the composite materials the growth characteristics of the *S. oneidensis* wildtype was compared with the *Δ cymA* strain, and a strain that was devoid of any outer membrane cytochrome *(Δ omc*). To this end, all cells were pre-cultured under oxic conditions at 30 °C in lysogeny broth (LB) medium. Cells were then transferred in LB medium or minimal medium (11.8 mM HEPES buffer, 1.27 mM $K_2HPO_4$, 0.73 mM $KH_2PO_4$, 2 mM $NaHCO_3$, 9 mM $(NH_4)_2SO_4$, 150 mM NaCl, 1 mM $MgSO_4$, 0.1 mM $CaCl_2$, 1 g $L^{-1}$ casein hydrolysate, 20 mM Na-D,L-lactate and trace elements (5 µM $CoCl_2$, 0.2 µM $CuSO_4$, 57 µM $H_3BO_3$, 5.4 µM $FeCl_2$, 1.3 µM $MnSO_4$, 67.2 µM $Na_2EDTA$, 3.9 µM $Na_2MoO_4$, 1.5 µM $Na_2SeO_4$, 5 µM $NiCl_2$, and 1 µM $ZnSO_4$) with an adjusted pH of 7.4) containing 50 µg $mL^{-1}$ kanamycin and cultivated under oxic conditions at 30 °C for 4 h. Cells were washed with freshly prepared medium and a suspension of *E. coli* or *S. oneidensis* strains with an optical density of 0.01 at 600 nm was prepared. After addition of 10 mM arabinose the suspension of *E. coli* or *Shewanella* strains was transferred into *S*-/*SC*-coated 96-well glass bottom plates. The growth of bacteria was monitored by fluorescence microscopy at 37 °C for *E. coli* and 30 °C for *Shewanella* strains, respectively.

To illustrate on-demand breakdown of the biohybrid materials and concomitant cell release, cultured materials grown for 24 h in LB medium were subjected to enzymatic cleavage. To this end, the biohybrid system was rinsed with PBS 3 times and digestion buffer (50 mM potassium acetate, 20 mM tris-acetate, 10 mM magnesium acetate, 100 µg/mL BSA, pH 7.4) 3 times. A solution of the restriction endonuclease BstEII-HF (10 U/mL in 300 µL digestion buffer) was added and incubated for 1 h at 37 °C. The cell release was monitored by confocal fluorescence microscopy. As a negative control, a solution containing SexAI was used under the same conditions. To investigate whether the release process led to viable cells, the released cells were re-suspended in fresh LB medium to an optical density of approx. 0.1 at 600 nm and their growth was monitored by recording the optical density at 600 nm of the cell suspension at 30 °C.

#### Scanning electron microscopy (SEM) analysis

After lyophilization, nanocomposite materials were coated with 4 nm of platinum using ion beam deposition and their morphology was characterized using a QUANTA 650-FEG scanning electron microscope (FEI Company) with an accelerating voltage of 5-10 kV.

Prior to resolving the *S. oneidensis* by SEM, *S. oneidensis* grown in the hydrated materials were fixed with PFA (4%, Polysciences) in DPBS at 4 ºC for 1 h, and freeze-dried after replacing DPBS with distilled water.

#### Fluorescence imaging analysis

Fluorescence micrographs of the materials and bacteria were recorded with an LSM 880 (Carl Zeiss) equipped with a temperature controlled container for cell growth. During live-cell imaging, the temperature of the container was held at 37 ºC for *E. coli* and 30 ºC for all *S. oneidensis* strains, respectively.

**Table S1.** List of DNA sequences.

| Name | Sequence (5’-3’) | Modification |
|---|---|---|
| aP1 | [AmC12]TCTAACTGCTGCGCCGCCGGGAAAATACTGTACGGTTAGA | 5’Amine C12 |
| P2 | TTTTTTTTTTTTTTTTTTTTTTTTTTTTTTTTTTTTTTTTCTAACTGCTGCGCCGCCGGGAAAATACTGTACGGTTAGA | - |
| T | [Phos]TTCCCGGCGGCGCAGCAGTTAGATGCTGCTGCAGCGATACGCGTATCGCTATGGGTAACCGTACGGTTACCCGCAGCAGCATCTAACCGTACAGTATT | 5’Phosphorylation |

**Table S2.** Dimensions of nanocomposite materials on interdigitated array electrodes.

| | Wet | Dry | |
|---|---|---|---|
| IDA Electrode | Area / $m^2$ | | Length / µm |
| 10 µm | 2,84E-07 | 1,42E-09 | 10 |
| 100 µm | 1,12E-05 | 5,60E-08 | 100 |
| 200 µm | 1,20E-05 | 6,00E-08 | 200 |

**Table S3**. Conductivity values of *S*, *SC* and *C* nanocomposite materials*

| | | 10 µm | | | | 100 µm | | | | 200 µm | | | |
|---|---|---|---|---|---|---|---|---|---|---|---|---|---|
| | | Dry | | Wet | | Dry | | Wet | | Dry | | Wet | |
| | | Mean | *STD* | Mean | *STD* | Mean | *STD* | Mean | *STD* | Mean | *STD* | Mean | *STD* |
| CP | *C* | 2,02E+01 | 3,48E+00 | 2,00E-05 | 9,83E-06 | 1,55E+00 | 2,50E-01 | 2,92E-06 | 1,26E-06 | 4,37E+01 | 1,27E+01 | 1,70E-05 | 7,84E-06 |
| | *SC* | 2,34E+01 | 4,94E+00 | 2,14E-05 | 8,70E-06 | 6,60E-01 | 8,10E-02 | 1,37E-05 | 1,19E-05 | 5,62E-01 | 3,96E-02 | 1,18E-05 | 6,14E-06 |
| | *S* | 7,86E-05 | -- | 2,51E-06 | 1,56E-06 | 1,14E-05 | 6,60E-06 | 2,53E-06 | 7,89E-07 | -- | -- | 8,38E-06 | 3,78E-06 |
| EIS | *C* | 2,23E+01 | 1,78E+01 | 3,56E-06 | 2,06E-06 | 2,63E+00 | 5,63E-01 | 3,82E-07 | 5,76E-08 | 3,12E+06 | 5,39E+06 | 3,19E-06 | 1,88E-06 |
| | *SC* | 2,83E+00 | 3,94E+00 | 6,02E-07 | 4,72E-07 | 8,06E-01 | 5,59E-01 | 3,59E-07 | 2,01E-07 | 4,63E-01 | 2,59E-01 | 1,49E-06 | 2,22E-07 |
| | *S* | 3,24E-05 | 4,77E-05 | 1,79E-06 | 9,10E-07 | 1,24E-07 | 1,43E-07 | 6,90E-07 | 5,25E-07 | 9,26E-08 | 1,13E-07 | 2,30E-06 | 8,18E-07 |

* Data was determined from triplicate chronopotentiometry (CP) and electrochemical impedance spectroscopy (EIS) measurements. Conductivity reported on 10, 100 and 200 µm electrode distances in hydrated and dehydrated states. Average values with standard deviation (S.D.) are reported for each sample. Missing data points for the dehydrated *S* materials are caused by failure to establish a steady state chronopotentiometry measurement.

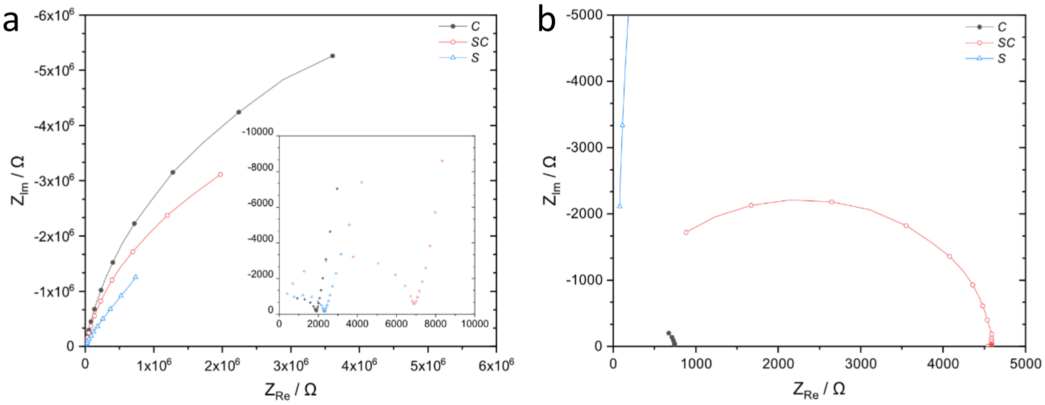

**Figure S1**. **Nyquist plots from electrochemical impedance measurements.** Nyquist plots from electrochemical impedance measurements of (**a**) hydrated and (**b**) dehydrated nanocomposite materials.

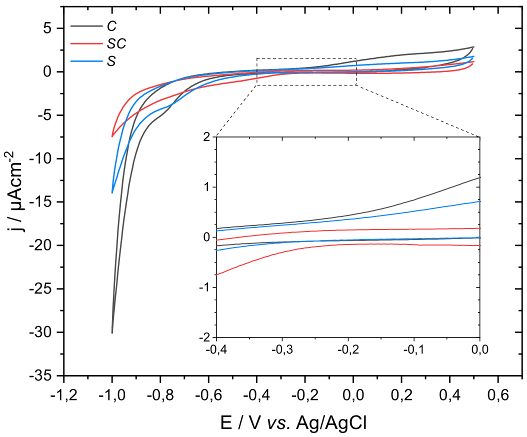

**Figure S2**. **CV analysis of nanocomposite materials without a redox mediator**. Inset shows the electrochemical activity in a potential window relevant for most bioelectrochemical systems.

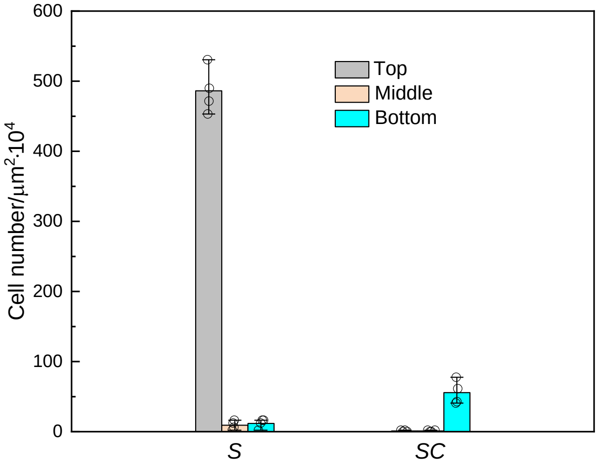

**Figure S3. Statistical analysis of *E. coli* grown in *S* and *SC* nanocomposite materials in LB medium.** After 24 h cultivation in LB medium, the cell numbers of *E. coli* grown in three different regions (i.e., top layer, middle layer and bottom layer) of *S* and *SC* nanocomposite materials was determined from fluorescence micrographs. Note that, *E. coli* can propagate well on top of the *S* nanocomposite materials, whereas almost no growth occurs in the *SC* nanocomposite materials (see also Figs. 3a, b in the main text).

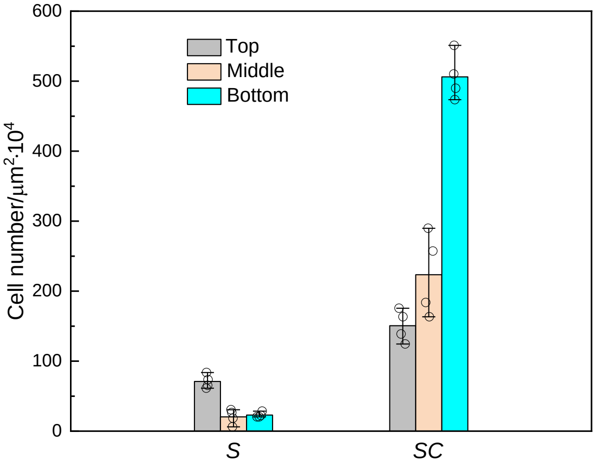

**Figure S4. Statistical analysis of *S. oneidensis* grown in *S* and *SC* nanocomposite materials in LB medium.** After 24 h cultivation in LB medium, the cell number of *S. oneidensis* grown in three different regions (i.e., top layer, middle layer and bottom layer) of the *S* and *SC* nanocomposite materials was determined from fluorescence micrographs. Note that *S. oneidensis* does not grow well in the *S* nanocomposite materials, whereas strong growth occurs in *SC* nanocomposite materials that contain CNT (see also Figs. 3c, d in main text).

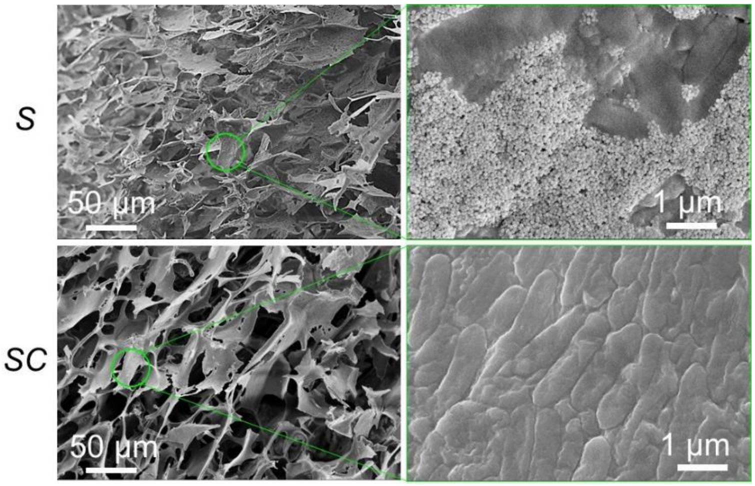

**Figure S5. SEM analysis of *S. oneidensis* grown in the nanocomposite materials.** Representative SEM images of *S. oneidensis* grown in *S* and *SC* nanocomposite materials in LB medium for 24 h, respectively. Higher magnification images in the right panels were acquired from the indicated green circular regions.

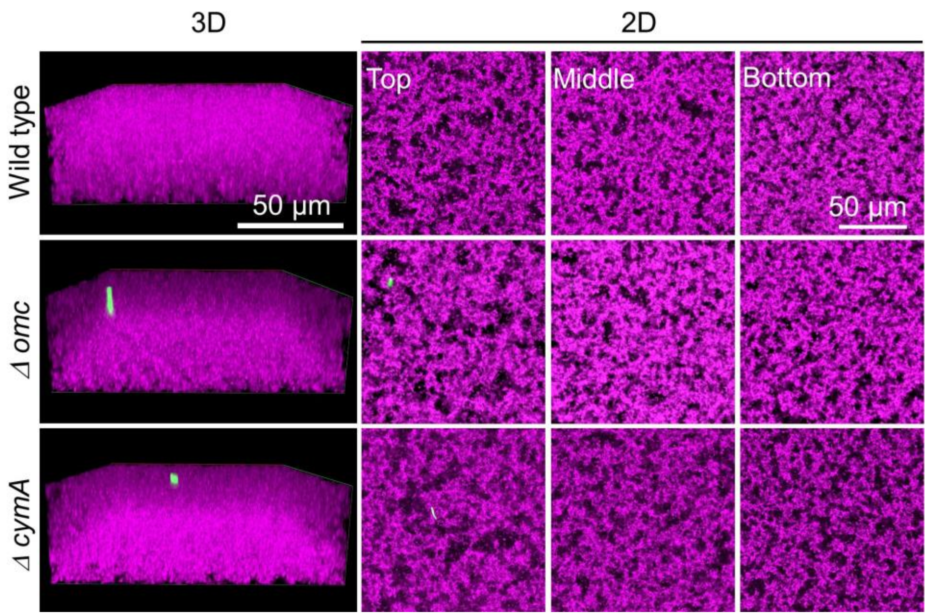

**Figure S6. Bacterial propagation in *SC* nanocomposite materials in minimal medium for 2 h.** Representative 3D and 2D fluorescence images of *S. oneidensis* strains (green) grown in *SC* nanocomposite materials (magenta) in minimal medium for 2 h. Note that, since it was intended to investigate the bacterial propagation in the nanocomposites, only few cells (0-1.1 cells/μm$^2$·10$^4$) were initially inoculated. The low starting densities are confirmed by the fluorescence microscopy images.

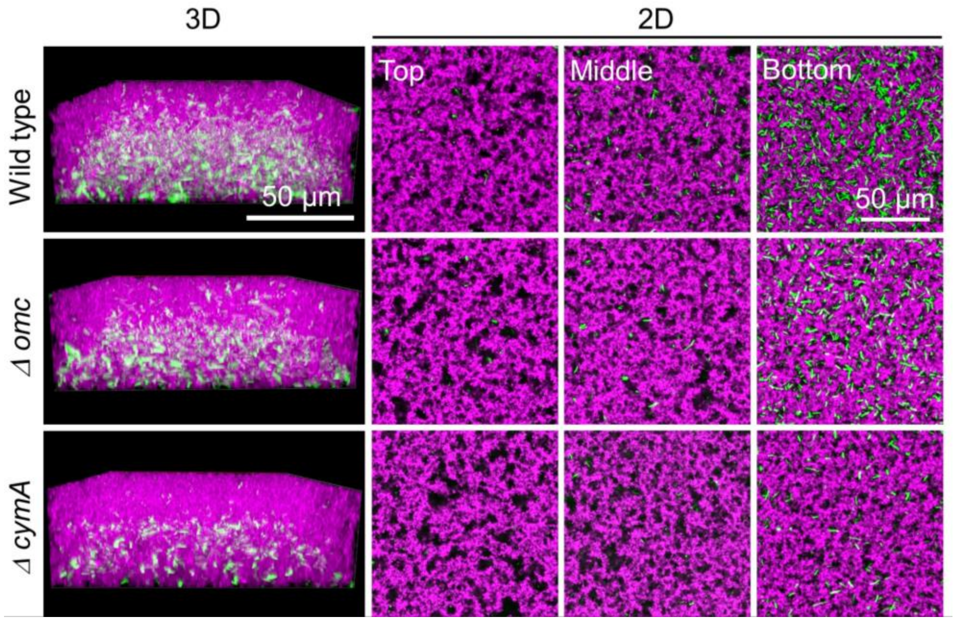

**Figure S7. Bacterial growth in *SC* nanocomposite materials in minimal medium after 48 h.** Representative 3D and 2D fluorescence images of the *S. oneidensis wild type*, *S. oneidensis Δ omc* and *S. oneidensis Δ cymA* (green) grown in *SC* nanocomposite materials (magenta) in minimal medium for 48 h.

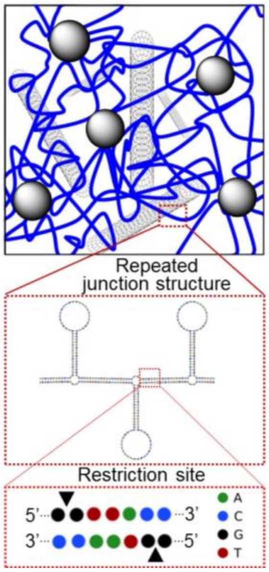

**Figure S8. Programming of restriction sites into nanocomposite materials.** Predicted secondary structures of RCA products obtained from circular template T, bearing stem-loop structures and double stranded regions that contain BstEII restriction sites in a repeated junction structure. Secondary structure of the RCA product structures were predicted by using the Nupack software.[3]

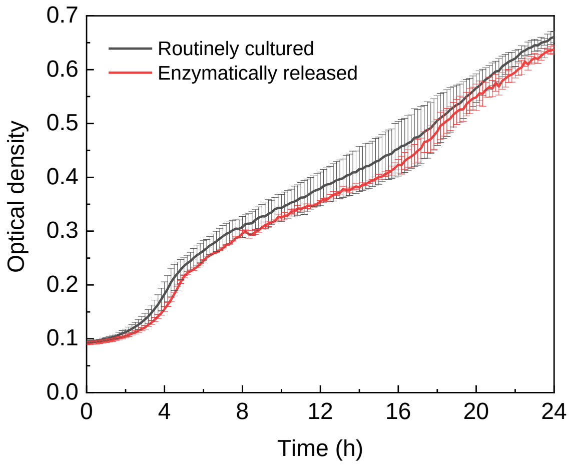

**Figure S9. Growth profile of the enzymatically released *S. oneidensis/SC* biohybrid material**. The collected *S. oneidensis* material (red curve) dispersed in LB medium and cultured for 24 h and the optical density was recorded at 600 nm. As a control, a routine cultivation was done under the same conditions with fresh *S. oneidensis* cells (black curve).

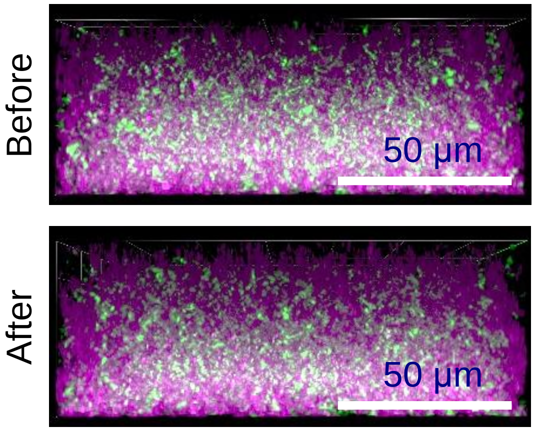

**Figure S10. Resistance of the *S. oneidensis/SC* biohybrid materials against the restriction enzyme SexAI.** Representative fluorescence images of the hybrid materials before and after treatment with the non-complementary SexAI restriction enzyme. Please note that the material system remains stable after treatment with this non-complementary endonuclease.
